# *Crowdsourcing*: Spatial clustering of low-affinity binding sites amplifies *in vivo* transcription factor occupancy

**DOI:** 10.1101/024398

**Authors:** Justin Malin, Daphne Ezer, Xiaoyan Ma, Steve Mount, Hiren Karathia, Seung Gu Park, Boris Adryan, Sridhar Hannenhalli

## Abstract

To predict *in vivo* occupancy of a transcription factor (TF), current models consider only the immediate genomic context of a putative binding site (BS) – impact of the site’s spatial chromatin context is not known. Using clusters of spatially proximal enhancers, or archipelagos, and DNase footprints to quantify TF occupancy, we report for the first time an emergent group-level effect on occupancy, whereby BS within an archipelago experience greater *in vivo* occupancy than comparable BS outside archipelagos, i.e. BS not in spatial proximity with other homotypic BS. This occupancy boost is tissue-specific and scales robustly with the total number of BS, or enhancers, for the TF in the archipelago. Interestingly, enhancers within an archipelago are non-uniformly impacted by the occupancy boost; specifically, archipelago enhancers that are enriched for BS corresponding to degenerate motifs exhibit the greatest occupancy boost, as well as the highest overall accessibility, evolutionary selection, and expression at neighboring gene loci. Strikingly, archipelago-wide activity scales with expression of TFs with degenerate, but not specific, motifs. We explain these results through biophysical modelling, which suggests that spatially proximal homotypic BS facilitate TF diffusion, and induce boosts in local TF concentration and occupancy. Together, we demonstrate for the first time cooperativity among genomically distal homotypic BS that is contingent upon their spatial proximity, consistent with a TF diffusion model. Through leveraging of three-dimensional chromatin structure and TF availability, weak archipelago binding sites *crowdsource* their occupancy as well as context specificity, with coordinated switch-like effect on overall archipelago activity.

## Introduction

Eukaryotic transcriptional regulation is critically mediated by the binding of specific transcription factors (TF) to their cognate DNA binding sites in the genome (Spitz and Furlong 2012). A TF’s *in vivo* DNA binding varies dramatically over developmental time and across tissues (Yáñez-Cuna et al. 2012; Plank and Dean 2014), and as such, a TF’s *in vitro* binding preference, or motif, does not accurately predict its *in vivo* binding (Yáñez-Cuna et al. 2012; Zinzen et al. 2009). Thus, a TF’s DNA binding motif suffers from being, both insufficiently informative to precisely specify binding in the large genomic substrate and insensitive to the *in vivo* environment, making it essential to characterize additional determinants of *in vivo* TF-DNA binding (Moses et al. 2004; Heinz et al. 2013).

Spatio-temporal variation in TF binding has been shown to be, in part, mediated by the local chromatin state of a binding site (BS) (Hesselberth et al. 2009). Recent work highlights three additional features of functional binding: (1) high GC content (White et al. 2013), (2) cooperative binding (Smith et al. 2013; Yáñez-Cuna et al. 2012) and (3) clusters of homotypic clusters of BS for a common TF, or HCTs (Gotea et al. 2010; Ezer et al. 2014a). These three features have been shown to be enriched in gene promoters and distal enhancers and to contribute to functional *in vivo* binding leading to transcriptional activation (Gotea et al. 2010; White et al. 2013; Arvey et al. 2012; Sharon et al. 2012). Still, most BS predicted by current models are not bound *in vivo* (Arvey et al. 2012; Slattery et al. 2014; Moses et al. 2004).

To date, research on determinants of functional TF binding, such as those above, have focused on putative BS and immediate flanking sequences. In parallel, the three-dimensional organization of the genome has emerged as an important mediator of transcriptional regulation, where spatial, not genomic, proximity is determinative (Fullwood et al. 2009; Ing-simmons et al. 2014; Babaei et al. 2015; Filippova et al. 2013). Chromatin looping events have been shown to combine at a higher organizational level, where they confer spatial proximity to functionally related genes and their distal regulatory regions (Fullwood et al. 2009; Li et al. 2012; Schwarzer and Spitz 2014; Fraser 2006; Lieberman-aiden et al. 2009). In vertebrates, for example, Hox genes, globin genes, and olfactory receptors, along with their distal enhancers, adopt a spatially clustered conformation, termed as ‘regulatory Archipelago’ (AP), as a prerequisite for robust transcriptional activation (Schoenfelder et al. 2010a; Vernimmen 2014; Montavon and Duboule 2012; Schwarzer and Spitz 2014; Markenscoff-Papadimitriou et al. 2014). Despite mounting evidence supporting functional criticality of chromatin interactions in context-specific transcriptional regulation, the potential impact of spatial clustering of BS on their individual TF occupancy has not been investigated. Recent findings that spatially clustered enhancers (we borrow the term ‘archipelago’ to refer to such spatially clustered enhancers) often share BS for the same TF, i.e., homotypic sites (Taher et al., 2013; Malin et al., 2013) make such enquiry even more compelling. Given the impact on functional binding of genomic homotypic clusters of BS (He et al. 2012b; Ezer et al. 2014a), we investigated the impact of spatial homotypic clusters of BS – that is, spatially clustered but genomically distant BS for a mutual TF.

In what follows, it's crucial to distinguish binding affinity of a TF for a BS, which is typically assessed *in vitro*, from TF occupancy at a BS, which is an *in vivo* state and depends on additional factors – most directly, TF concentration (Foat et al. 2006). Importantly, TF concentration and, hence, TF occupancy, may be non-uniformly spatially distributed in the nucleus (Schoenfelder et al. 2010a; Chakalova and Fraser 2010). Indeed, as described by facilitated TF diffusion, BS for a common TF in a HCT act together to briefly 'trap' a TF into diffusing back and forth amongst themselves along the chromatin (Brackley et al. 2012; Ezer et al. 2014a, 2014b), resulting in higher-than-expected occupancy in the HCT.

Here, based on clusters of spatially proximal enhancers, or *APs* (Malin et al 2013, Sheffield et al 2013), and using nucleotide-resolution DNase footprints to quantify context-specific *in vivo* TF occupancy data (Neph et al. 2012), we demonstrate a strong group-level effect on TF occupancy whereby individual BS within an AP experience greater *in vivo* occupancy than their counterparts outside APs, i.e., enhancers that are not in spatial proximity with other enhancers. We refer to the differential occupancy as *‘occupancy boost’*. Strikingly, occupancy boost for a TF in an AP scales robustly with both the number of putative BS per AP enhancer, i.e., homotypicity, as well as the number of accessible enhancers in the AP. TFs with degenerate motifs, which are expected to have abundant putative BS, are consistently among the TFs subject to the greatest occupancy boosts; in large APs, mean boosts for homotypic BS corresponding to highly degenerate motifs are between 2 and 3-fold. Based on these results, we propose that *in vivo* occupancy at particular BS in an AP is amplified by the presence of homotypic BS in spatial proximity, i.e., BS ‘*crowdsource*’ their own occupancy boost along with other homotypic BS in their spatial proximity. We extend the aforementioned biophysical model of facilitated diffusion of TFs within one HCT to show that the observed occupancy boost can result from TFs briefly ‘trapped’, or diffusing, among multiple spatially proximal HCTs. Among AP enhancers, we find that specifically those enriched in degenerate motifs (‘enriched enhancers’) are most affected when compared with AP enhancer depleted for degenerate motifs and comparable non-AP enhancers; they exhibit several-fold greater boost in activity, their neighboring genes exhibit several-fold greater expression and, consistent with functional significance, they exhibit a greater evolutionary conservation. Tellingly, mean AP-wide chromatin accessibility scales with the expression of cognate TFs with degenerate – but, not specific – motifs. Combined with other results, this points to a role for crowdsourcing in: (i) initiating a positive feedback loop whereby greater TF occupancy at enriched enhancer BS increases the overall accessibility at these enhancers, thus facilitating further occupancy; (ii) endowing enriched enhancers with switch-like behavior, activating them in specifically those tissues where chromatin structure and TF availability together result in sufficient occupancy boost

## Results

### Analysis overview

Our analysis is based on previously identified enhancer clusters (Malin et al. 2013) comprising ∼1600 enhancers in 40 clusters. Enhancers were clustered based on correlated DNase hypersensitivity (DHS) profiles across 37 cell lines (representing 82 cell lines). Enhancers in the same cluster were shown to (i) have functionally related gene neighbors with correlated expression, indicative of coordinated regulation, (ii) share BS for several TFs, and (iii) be spatially proximal to one another. We will refer to such enhancer clusters as 'archipelagos' (APs) borrowing from (Spitz and Furlong 2012). We refined the APs identified in (Malin et al. 2013) to ensure tight spatial proximity among AP enhancers (see Methods). Note that properties (ii) and (iii) above together imply a higher spatial density of homotypic BS within an AP, particularly for TFs with abundant putative BS (Fig. 1A); this generally corresponds to TFs with degenerate motifs.

**Figure 1.**
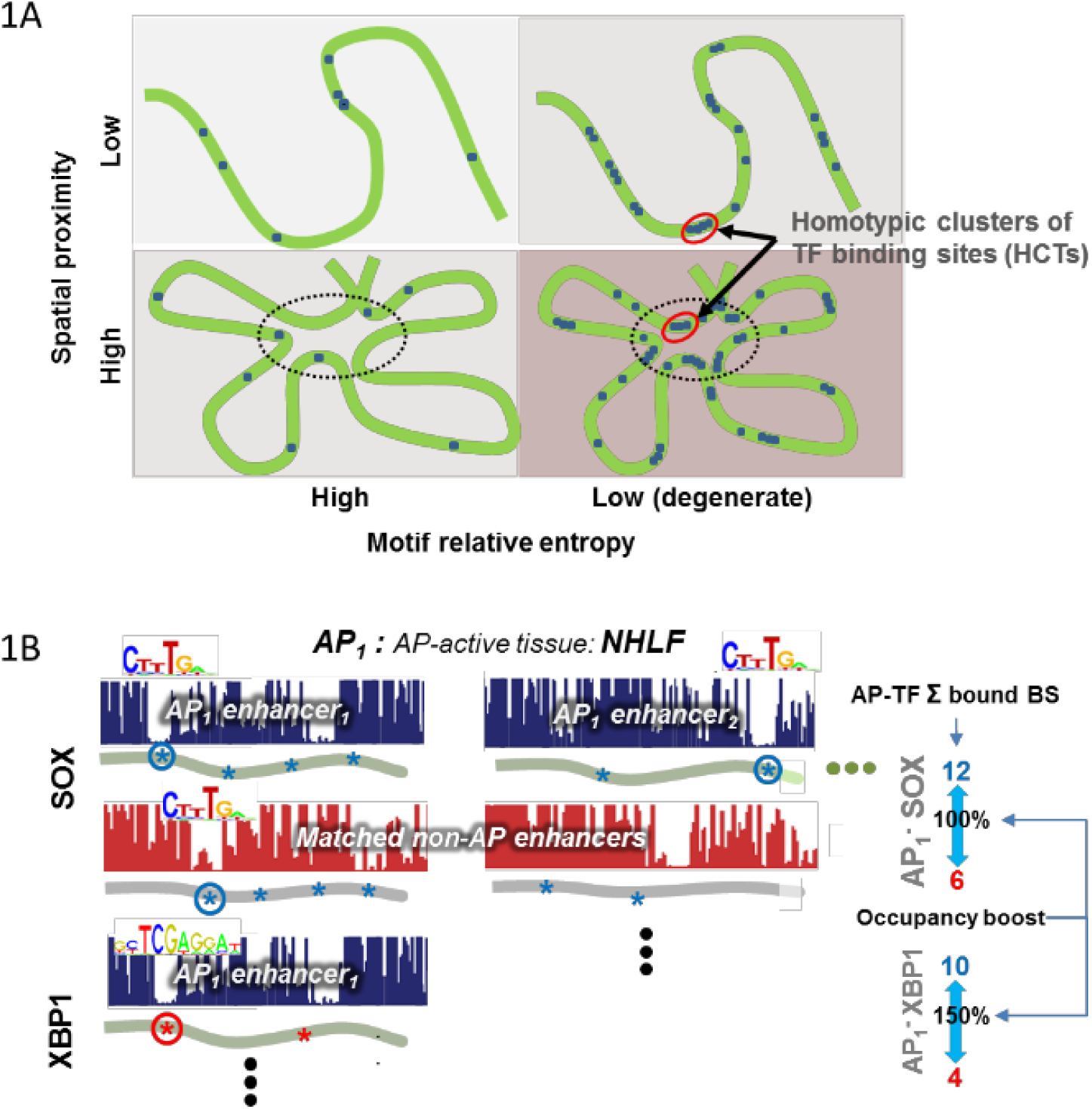
Expectation and testing of differential occupancy. **(A)** The combination of spatial proximity and genomic homotypic clusters of TFBS produce high homotypic TF BS concentration. As illustrated, low-RE (degenerate) motif BS have a higher expected frequency in the genome than high-RE motif BS, including more frequent HCTs. In a spatially proximal chromatin context, effective homotypic BS concentrations are particularly elevated for low-RE motif BS. This effect is further accentuated in archipelagos of enhancers, which have been shown to be enriched for HCTs for shared TFs. High effective homotypic BS concentration is likely a pre-requisite for the crowdsourcing effect. Large ovals denote archipelagos of functionally related enhancers and target genes. Darkness of background color approximates the maximum expected homotypic BS concentration. Not drawn to scale. Green: DNA. Black: BS. BS=binding site; RE=relative entropy; HCT = homotypic (genomic) cluster of TFBS. **(B)** Calculating differential TF occupancy boost based on curated digital DNase footprint data. Shown is the procedure for calculating occupancy boost for each (AP, TF) pair. For each enhancer in an AP, and each TF with one or more putative BS in the enhancer, a non-AP enhancer is chosen (with replacement) after controlling for mean enhancer-wide chromatin accessibility (DHS) in the AP’s most active tissue, and for the number of putative BS. For each TF-AP pair, then, occupancy boost is calculated as the percent difference in the number of putatively bound BS, where binding is determined in a binary manner: 1, if a curated footprint tightly overlaps a given motif instance, 0, otherwise. If multiple TF motifs tightly overlap a given footprint, conservatively, all are classified as bound. Putative BS are indicated by a ‘1’, or ‘2’, respectively, for example TFs SOX and XBP1. A circle around a BS signifies it is imputed as bound by its cognate TF. Note that the toy calculation of occupancy boost does not correspond to the data displayed. AP = archipelago; TF = transcription factor, BS = binding site. DNase digital footprint scans from Neph et al 2012.

Previous study suggests that genomic clustering of homotypic BS can boost the cognate TF’s occupancy at individual BS in the cluster (Brackley et al., 2012; Ezer et al., 2014b). By extending the notion of genomic clusters of homotypic BS to *spatial* clusters of homotypic BS, here we hypothesize that analogously, mediated by conformational changes in chromatin organization, such spatial BS cluster may boost binding occupancy for a cognate TF at individual BS, with potential downstream functional impacts. To test this hypothesis, we ***(i)*** contrasted *in vivo* TF binding occupancy in AP enhancers to that in a stringently controlled set of *‘non-AP’* enhancers in the same tissue, ***(ii)*** assessed functional impact of occupancy boost at AP enhancers – particularly those enriched for degenerate motif sites – and their putative gene targets, ***(iii)*** characterized context-specificity of the occupancy boost, and ***(iv)*** tested context-specific TF availability as an upstream driver of the occupancy boost.

In the following analyses, *in vivo* occupancy was estimated for each putative BS using, alternatively, high-resolution curated cell type-specific DNase footprint data (Neph et al. 2012) or ChIP-Seq (www.encodeproject.org/ENCODE) (see Methods). For additional validation, key tests were repeated using an alternative set of previously published APs (Sheffield et al. 2013). *In vivo* occupancy of a TF and other functional analyses were primarily performed in each AP’s most active cell line out of 34 examined– its so-called ‘AP-active’ tissue (see Methods), which offered 730K BS for analysis and 1.8m more in the alternative AP dataset. Results in the AP-active tissue were then contrasted with those in ‘AP-inactive’ cell lines. With respect to background for BS-level analyses, for a given TF, each AP enhancer was matched to a non-AP enhancer controlling for chromatin accessibility and the number and type of putative BS (Methods).

### Occupancy boost at AP BS increases with homotypic BS density within AP, supporting crowdsourcing of *in vivo* TF occupancy

A putative BS for a TF is deemed bound *in vivo* by the TF if the BS overlaps DNase footprint based on stringent criteria (Fig. 1B and Methods). For a collection of homotypic BS, occupancy is quantified as the proportion that are bound *in vivo*. We first found that the *in vivo* occupancy across all BS in all APs in their respective AP-active cell lines (Methods) is significantly higher than the occupancy across matched non-AP enhancers (Wilcoxon test p-value =2.5E-7), albeit with modest effect size (odds ratio of occupancies = 1.21). However, according to our central hypothesis we expect AP occupancy to be pronounced primarily for the subset of TFs with large numbers of BS in spatial proximity in an AP (Fig. 1A). We therefore examined such TFs explicitly.

For a given TF in one AP, we calculated the TF’s *coverage* as the total number of cognate BS in the AP, and calculated its *occupancy boost* as the ratio of occupancy at these BS to the occupancy at a matched set of non-AP enhancer BS (Methods). Such TF-AP pairs are the fundamental unit at which the crowdsourcing hypothesis predicts occupancy will be impacted. Because background levels of BS occupancy in the genome are generally low (3-4%), the occupancy is zero in both AP and control non-AP enhancer sets for a majority (65%) of the 25K TF-AP pairs; these pairs were excluded for this analysis. Of the remaining TF-AP pairs, 3.6k have non-zero occupancy in both AP and non-AP, encompassing ∼95K enhancer-TF pairs and ∼205K BS (we call this the *reciprocal* set), and additional 5k TF-AP have non-zero occupancy in either AP or in matched non-AP BS (*non-reciprocal* set). We analyze the two sets of TF-AP pairs separately.

We stratified the reciprocally occupied TF-APs into 8 bins with exponentially increasing coverage cutoffs and calculated the overall occupancy boost for each bin as the mean occupancy boost among member TF-APs. A 95% confidence interval for each bin’s mean was estimated using a bootstrap. As shown in Fig. 2A, the occupancy boost robustly increases with the TF coverage in the AP. Specifically, we found a substantial difference in occupancy boost between TF-APs with the highest and lowest 50% coverage (mean of 77.7 % versus 2.1 %; Wilcoxon p-value = 1.4e-5). This trend also holds when coverage was alternatively quantified as the number of enhancers in an AP with at least one BS for the TF (Fig. 2B), suggesting that the boost is not due to disproportionate contribution from a few enhancers, but instead relies on widely dispersed BS across the AP’s enhancers. Interestingly, the boosts for high coverage TF-APs increase when the digital footprint binding criterion for assessing occupancy is made more stringent (Supplemental Fig. S1A).

**Figure 2.**
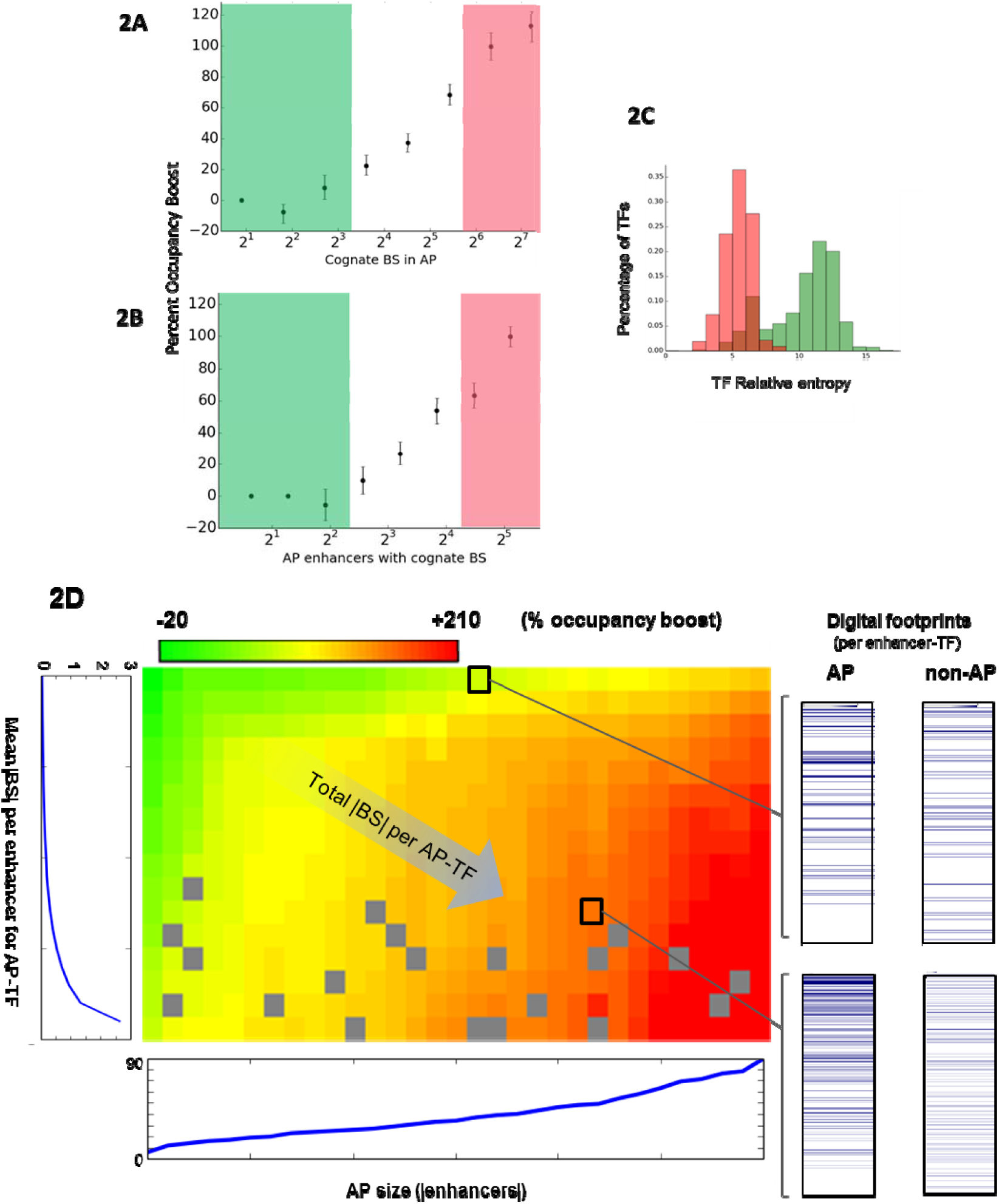
Differential AP occupancy ‘boost’ scales with TF coverage in the AP. TF-AP combinations were sorted on the basis of coverage and mean occupancy boost was determined for each group of TF-APs, where occupancy boost refers to differential occupancy in AP and non-AP enhancers matched 1-to-1 for the TF’s motif signature (the number and type of motifs) in a given enhancer, as well as for mean DHS across the AP. Occupancy was calculated based on the overlap of curated DNase digital footprints (Neph et al 2012) with high-confidence TRANSFAC motif instances. TF-APs and their non-AP counterparts were included in this analysis only if they both had non-zero occupancy (See also Figures S1, S2, S3). Coverage was calculated alternatively as: **(A)** the number of cognate BS for a given TF in a given AP or **(B)** the number of enhancers in the AP with one or more motif instances cognate to the TF. **(C)** There is a strong inverse relationship between TF-AP coverage and TF relative entropy, as expected. TF-APs with the top (salmon) and bottom (green) 20% coverage were mapped to the TFs’ RE values. Colors in 2A and 2B illustrative and do not coincide with these coverage ranges. AP: archipelago, RE = relative entropy. **(D)** Mean occupancy boost versus coverage that has been decomposed along two axes. Each TF-AP pair was binned based on the number of enhancers in an AP (column) and the mean number of BS per AP enhancer (row). Plots to the left of and below the heatplot show mean boost for each row and column, respectively. Red (green) heatmap cells indicate high (low) percentage occupancy boost after Lowess smoothing. Grey cells indicate no data. In the right panel, for all TF-AP pairs in the selected heatmap cell, significant digital DNase hypersensitivity footprints in member AP and matched non-AP enhancers are shown, where the numbers of BS for AP and non-AP enhancer-TF pairs are identical; a blue line indicates a significant footprint overlapping a putative BS. Enhancers are sorted from bottom to top in order of increasing chromatin accessibility.

Abundance of a TF’s cognate BS is strongly correlated with its motif degeneracy, as quantified by the motif’s relative entropy (RE) (Fig. 2C) – low RE is identified with high degeneracy (Hannenhalli 2008). Given this association, we also directly assessed the relationship between TF motif RE and occupancy and found very similar trends (Supplemental Fig. S1B). Interestingly, the steepest declines in occupancy boost and in the distribution of TF RE echo one another, as shown, in that both occur at RE = 7. When, in an independent set of 11 tissues, we instead used ChIP-Seq to infer occupancy the trend, again, remained very similar (Supplemental Fig. S1C). Taken together, the above analyses strongly suggest that binding sites for high coverage TFs experience a substantial occupancy boost in AP enhancers relative to BS in comparable non-AP enhancers.

The overall TF coverage is affected by both the mean number of BS per AP enhancer ('homotypicity') and the number of enhancers per AP ('AP size'). Next, we assessed the relative contributions of these two constituents of coverage on the occupancy boost. As shown in Fig. 2D, for the reciprocal set, AP size and homotypicity independently and robustly impact the magnitude of occupancy boost (p-value = 4.2e-6). A similar analysis on 5K non-reciprocal TF-AP pairs shows a similar and significant trend (Supplemental Fig. S1D; p-value 8.1E-5). Our overall conclusion does not change when we used an independent set of 450 AP enhancer clusters reported in (Sheffield et al. 2013) based on 1.8 million BS (supplementary Note 1) (Figs. S2A, S2B). Consistent with the expected importance of spatial proximity, as the Hi-C based screen for pairwise distance between fellow AP member enhancers became more stringent, the trend improved (Supplementary Note 1).

The observed link between spatial BS abundance and occupancy, we reasoned, may partly be mediated by cooperativity among the bound TFs within an AP (Martinez and Rao 2012). However, as explained in supplementary Note 2, we found that even though heterodimerzing TFs exhibit a somewhat (but significantly) larger boost than other TFs, this difference can largely be explained by their greater BS coverage (Supplemental Fig. S3. Taken together, these results strongly suggest that AP TF’s occupancy at a specific BS in an AP is ‘*crowdsource*d’ by the collective of its putative BS in the AP.

### TF occupancy boost in spatial clusters of BS is consistent with a facilitated-diffusion model

Previous studies have shown that a facilitated diffusion model can explain the greater occupancy in homotypic clusters of BS (Brackley et al. 2012). Here we simulated an extended version of the biophysical model for HCTs in isolation to determine whether the crowdsourcing effect is sufficient to explain the observed AP-mediated occupancy boost. All details pertaining to the model, algorithms, and results are provided in Supplemental File ‘Facilitated_Diffusion_Model’. First, in a simple scenario, we investigated specifically how occupancy is affected by (i) the spatial distance between enhancers within an AP cluster, (ii) the genomic distance between binding sites within an AP enhancer, and (iii) the number of binding sites within each enhancer. Then, we investigated various 3D organizations of homotypic binding sites that would closely represent the crowding of low-RE binding sites in AP clusters *in vivo*. Our simulation results suggest that both the presence of homotypic clusters and inter-strand jumping between enhancers within an AP can increase the average TF occupancy. For instance, in the case of four enhancers containing pairs of homotypic binding sites, as shown in Fig. 3B, there was a 60% to 170% increase in TF occupancy when the enhancers were part of AP clusters (in which enhancers are 100nm to 200nm apart, as compared to being 10000 nm apart, by default), and in the case of eight enhancers containing pairs of homotypic binding sites there was an 118% to 277% increase occupancy (Fig. 3D), which suggests that the crowdsourcing effect is a biophysically sound strategy for increasing local TF occupancy in APs at a biologically meaningful scale.

**Figure 3.**
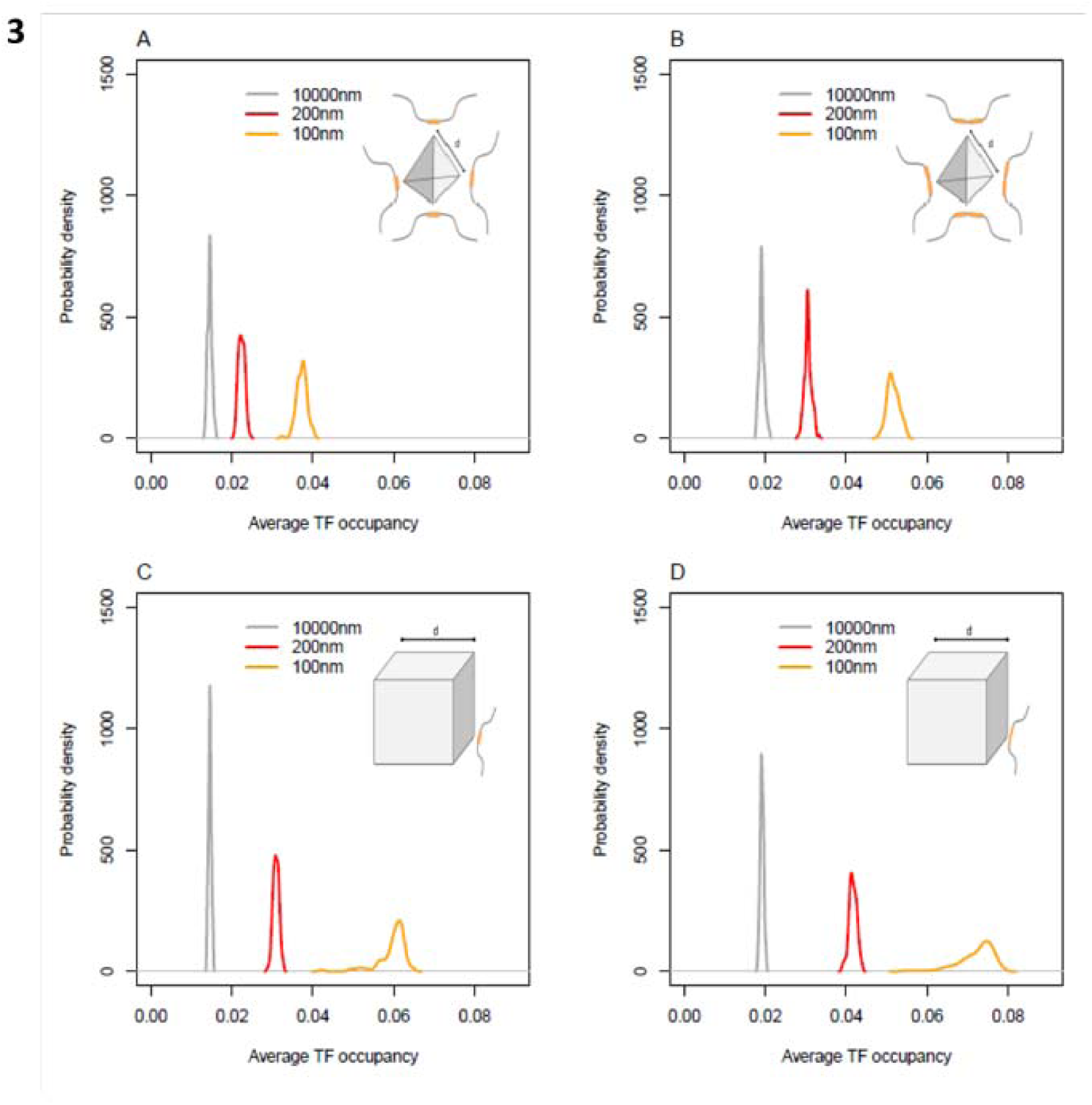
Biophysically modeling crowdsourcing effect. TF diffusion was simulated for four geometric arrangements of binding sites, and the probability density functions of TF occupancy are shown. The TF occupancy is defined as the average probability that each site is bound. The four simulated scenarios are: a tetrahedron with **(A)** one binding site or **(B)** a pair of binding sites in each corner, which contain 4 or 8 binding sites, respectively; a cube with **(C)** a single binding site or **(D)** a pair of binding sites in each corner, which contain 8 and 16 binding sites respectively. For an additional figure and details on the simulation, see Supplemental Methods.

### AP enhancers that are enriched for degenerate motifs have greater occupancy boost and higher downstream functional impact

Previous studies have shown a trend for AP enhancers to share homotypic BS (Malin et al 2013, Sheffield et al 2013). We reasoned that this could result if AP enhancers were enriched for degenerate, i.e., motifs with low RE, which are expected to be generally abundant, thus increasing the chance of being shared among AP enhancers. We found this to be true (Fig. 4D). Furthermore, our empirical results above and the biophysical model suggest that homotypicity within an enhancer as well as spatial clustering of such enhancers can increase TF occupancy at biologically meaningful scales. Together, these led us to hypothesize that the particular AP enhancers that are enriched for low-RE (degenerate) motif BS may be especially affected by crowdsourcing. In the following analyses, we employ multiple thresholds for RE to define ‘low’ RE motifs.

**Figure 4.**
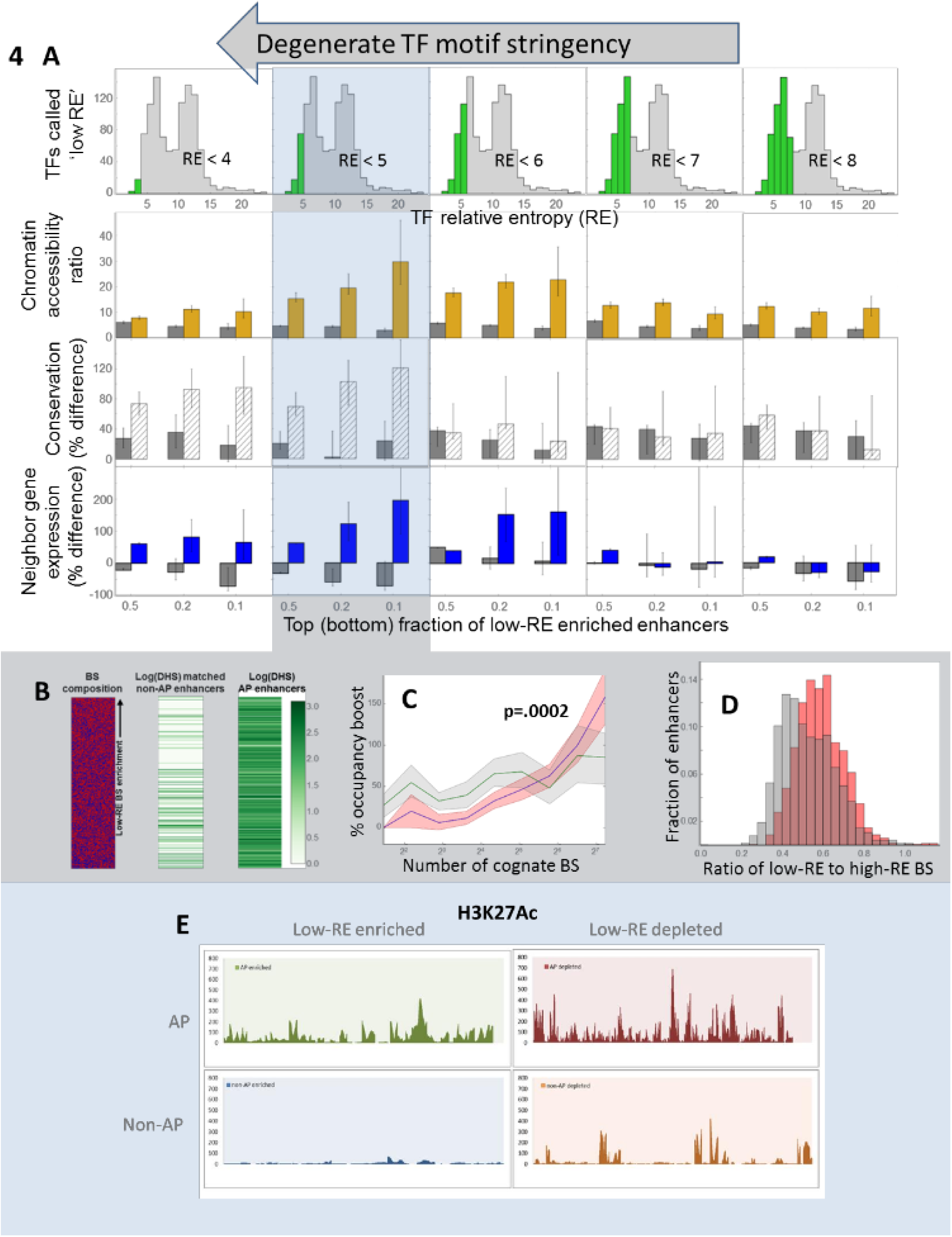
Enhancers enriched for degenerate BS are more functional than expected. **(A)** Enhancers enriched and depleted for low-RE BS were compared in terms of DNase hypersensitivity (row 2), evolutionary constraint (row 3), and neighbor gene expression (row 4). Readouts on y-axes indicate values normalized against carefully matched non-AP enhancers for the given RE cutoff (column). Within each plot, the 10%, 20%, and 50% (x-axis) most enriched enhancers are indicated in non-grey, while the most depleted enhancers are shown in grey. The histograms in the top row indicate the fraction (green) of all TFs deemed low-RE for the purpose of calculating each enhancer’s low-RE BS enrichment. Figures **(B-E)** are for RE cutoff = 5. **(B)** Chromatin accessibility of AP and matched non-AP enhancers, sorted by low-RE motif enrichment. Aligned heatplots display one enhancer per row (most enriched at top). (Left) heatplot in which red signifies a low-RE motif and blue a high-RE motif. Low-RE motif enrichment based on Fisher exact test against a background that included all non-AP enhancers. For visualization purposes, enhancer lengths and BS lengths standardized. (left, right). Log of non-AP and AP enhancer DHS, respectively. (See also Figure S4). **(C)** Percentage occupancy boost is shown as a function of coverage for AP enhancers with the highest 20% enrichment (blue line) and the highest 20% depletion (green line) for low-RE BS, along with their 95% confidence intervals. Coverage for a given TF-AP pair was calculated as the number of cognate BS in in the AP among enriched (depleted) enhancers only. A p-value is given for a Wilcoxon test comparing boosts among TF-APs with top 20% coverage. **(D)** Ratio of low-RE to high-RE motifs in AP enhancers vs. non-AP enhancers. AP and non-AP enhancers were matched one-to-one for DHS in each AP enhancer’s most active tissue. Putative BS were identified based on 95 percentile motif match threshold. The x-axis shows the ratio of low-RE to high-RE motif sites in each enhancer. Y-axis shows percentage of enhancers analyzed. Red: AP enhancers, Gray: non-AP enhancers. All confidence intervals were calculated using an appropriate booststrap method. AP: archipelago, RE: relative entropy, BS: binding site, TF: transcription factor. **(E)** Acetylation levels in enriched vs. depleted enhancers. Juxtaposed views of H3K27Ac ChIP-Seq in HUVEC are shown for 40 (100) AP enhancers in the top row that are in an AP that is active and in the top 10% for enrichment (depletion). Shown in the bottom row are views for matched non-AP enhancers.

Activity of an enhancer, and ultimately, activation of its target gene, is associated with the amount of TF binding in the enhancer (Smith et al. 2013; Fisher et al. 2012). Hence, given the occupancy boost at low-RE TF sites in AP enhancers, we hypothesized that the enhancers that are enriched for low-RE TF sites will have the largest overall increase in activity, and by proxy, increase in expression of their putative target genes. For each AP enhancer, assuming its closest gene neighbor to be its putative target (Djebali et al. 2012), we calculated its ‘expression boost’, similar to occupancy boost above, as the relative change in expression of the gene relative to the target gene of the control non-AP enhancer (Methods). We then compared expression boost for AP enhancers enriched for low-RE motif BS (*enriched* enhancers) to that for AP enhancers depleted in low-RE sites (*depleted* enhancers) (Methods). As shown in Fig. 4, the putative target genes of enriched enhancers have much greater expression in AP-active tissues than their non-AP counterparts, while the depleted enhancers do not. Moreover, as the degree of enrichment increases from top 50% to top 10%, the relative expression boost increases from 62% to 196% (for low-RE cutoff of 5); The difference in AP and non-AP neighbor gene expression for depleted enhancers is nearly as stark – but in the opposite direction: as the degree of enrichment increases from top 50% to top 10%, the drop in AP expression relative to non-AP grows from -32% to -72% (i.e., non-AP expression is 3.5-fold higher). GC content differences cannot explain either trend, as GC content in non-AP enriched (depleted) enhancers is <10% (15%) higher than in corresponding AP enhancers. Also, notably, the highest expression boost (Figure 4A, row4) and the highest occupancy boost (Fig. S1B) occur at similar low-RE cutoffs, thus strengthening the link between the two; at higher low-RE cutoff (being more permissive) the observed effect weakens and eventually disappears.

As a complementary test (Supplementary Note 3), we compared TF binding patterns between enhancers within 50Kb of highly expressed genes and enhancers within 50kb of lowly expressed genes. Consistent with above, we found that the low-RE motif usage (defined as the fraction of bound BS that are low-RE) was 1.8 to 3.0 times higher in AP enhancers near highly expressed genes than in those near lowly expressed genes. As a negative control, we did not observe this pattern for matched non-AP enhancers. Taken together, these results suggest that low-RE binding specifically at enriched AP enhancers have a significant impact on downstream gene expression. As an additional ascertainment of the functional importance of enriched AP enhancers, we found such enhancers to be up to 120% (90%) more evolutionarily conserved at RE cutoff of 5 (4) (using 20 species PhastCons score (Siepel et al. 2005)) than matched non-AP enhancers; indeed, the greater their enrichment, the greater the evolutionary constraint we observed (Fig. 4A, row 3). Depleted AP enhancers, by contrast, were at most 40% more conserved than their non-AP counterparts.

TF binding and chromatin accessibility are intimately connected; higher accessibility typically leads to higher occupancy, while TF binding can help displace a nucleosome and increase accessibility (Teif and Rippe 2012). Therefore, we assessed whether enriched enhancers exhibit a greater boost (relative to matched non-AP enhancers) in overall accessibility compared with depleted enhancers. For this analysis, we normalized AP enhancer accessibility by that of non-AP enhancers matched one-to-one with AP enhancers for numbers of low and not-low RE BS, as described previously, except DHS was explicitly left uncontrolled. As shown in Fig. 4A (row 2) and Fig. 4B, at RE threshold of 5, the most enriched enhancers exhibit ∼10-fold greater DHS boost (with respect to matched non-AP enhancers) than do depleted enhancers; as expected, the trend weakens for higher RE thresholds. To further resolve the effect of BS RE on enhancer accessibility, we tracked changes in accessibility as we increased the number of low-RE (high-RE) sites, while holding relatively constant the number of high-RE (low-RE) sites. As shown in Supplemental Fig. S4, increasing the number of low-RE BS (for relatively constant high-RE BS count) has a substantial positive impact on enhancer’s accessibility – especially when the number of high-RE BS is low – while increasing the number of high-RE BS (for a relatively constant low-RE BS count) has an insignificant or negative impact. In addition, we found that histone acetylation levels (H3K27Ac), which are associated with active enhancers, had a ratio between enriched and depleted AP enhancers 3-fold higher than the same ratio in non-AP enhancers (Fig. 4E).

Finally, we observed occupancy boosts in enriched enhancers that, unexpectedly, were up to two-fold higher than in depleted enhancers, often in the same AP and for the same levels of coverage (22o% vs. 110% at low-RE cutoff of 4 (not shown); 93% vs 39% in alternative APs (Sheffield et al 2013) (Supplementary Note 4; Fig. 4C, Supplemental Fig. S2C). To account for these unexpectedly high boosts, we address the potential for higher-order interactions in enriched enhancers among BS for distinct TFs with degenerate motifs (see Discussion)

These results – the relatively higher occupancy boosts, chromatin accessibility, downstream gene expression, and evolutionary constraint in enriched enhancers, along with greater prevalence of enriched enhancers among AP than non-AP enhancers – strongly suggest a hitherto unreported special functional relevance of AP enhancers that are enriched for low-RE binding sites.

### Cell type-specificity of AP enhancer occupancy boost and activity

Given the link between occupancy boost and spatial clustering of BS, and given the context-specificity of spatial proximity (Ay et al. 2014), we expect the occupancy boost to exhibit cell type specificity. In addition to identifying the cell type where an AP is deemed active (as employed in analyses thus far), we also identified the cell types where an AP is deemed inactive, namely those where less than 40% of the AP enhancers were DNase I hypersensitive. To offset the paucity of bound sites in inactive tissues, all qualifying inactive tissues for each AP were pooled. We found that for the TF-AP pairs in the highest coverage bin, occupancy boost dropped from ∼112% in its AP-active cell type down to 38% in inactive cell types (Fig. 5A). In addition, we estimated tissue specificity of each TF as the cross-tissue dynamic range of its occupancy, defined as the ratio of its occupancy in AP-active tissue(s) to that in AP-inactive tissues, calculated over the identical AP BS. Notably, this provides evidence of the occupancy boost’s tight association with coverage without the need for non-AP occupancy as a baseline (Methods). After controlling for DHS across coverage bins, we find that the TF-APs with top 10% coverage display 135% greater occupancy in active relative to inactive tissues, while in the matched non-AP context it is 38% (Fig. 5B). Even larger differentials between AP and non-AP contexts were observed for their respective ratios of non-reciprocal binding in active and inactive tissues (Supplemental Fig. S5A). Interestingly, we found that high coverage TF-AP pairs for heterodimerizing TFs exhibit substantially higher specificity than other TFs (225% vs. 140%) (Supplemental Fig. S5B), particularly TFs in MADS and bZIP domain families, suggesting an augmented level of cooperative binding in APs. This, we suspect, is due to the relatively binary nature of cooperative binding: in response to small increments in TF concentrations, heterodimers exhibit disproportionately large changes in occupancy (Giorgetti et al 2010).

**Figure 5.**
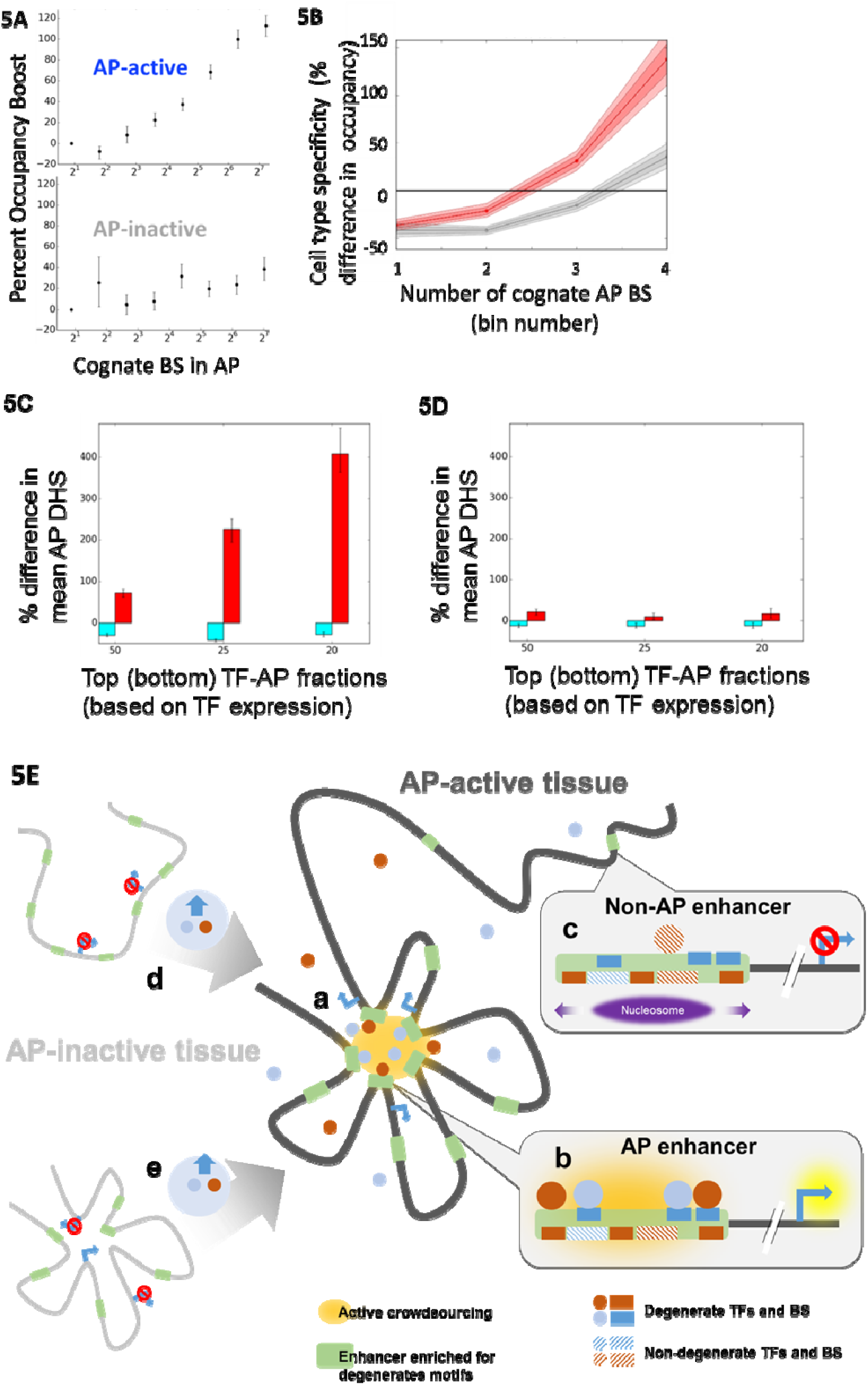
Mean AP accessibility scales with context-specific availability of TFs with degenerate motifs and not the TFs with specific motifs. **(A)** Occupancy boost in cell types with reduced AP activity. Occupancy was computed as a function of coverage in ‘inactive’ cell types – those in which fewer than 40% of the AP’s enhancers were DNase hypersensitive (bottom). For comparison, the plot for active cell types from 2A is reproduced (top) **(B)** Tissue specificity of occupancy for TF-AP and matched non-TF-AP pairs as a function of TF-AP coverage. Dynamic range (y-axis) for occupancy was calculated for each TF-AP pair as the percentage difference between mean occupancy in the AP’s most active and inactive cell types. Identical BS in active and inactive cell types were tested. Serving as a control, dynamic range was also computed for non-TF-APs that were matched to TF-APs. A TF-AP was required to have non-zero occupancy in both inactive and active cell types. Only TF-APs shared in AP and non-AP contexts were used for analysis. Shown are results for TF-APs and for non-TF-APs each sorted into 4 bins with exponentially increasing coverage cutoffs. Red: AP, Gray: non-AP. 90% and 99% confidence intervals are shown with variable hue. AP-wide DHS as a function of context-specific TF availability. (See also Supplemental Fig. S5) **(C)** For each TF-AP, tissue specific DHS was compared across each of 15 tissues for which there was RNA-Seq data available. (TF, AP, tissue) triplets were segregated into lowest-20%-coverage (cyan) and highest-20%-coverage (red) classes based on TF-AP, and then further subdivided into low and high expression based on tissue-specific TF expression. Bar height indicates the percentage increase in DHS level associated with an increase in TF expression from bottom <x> to top <x> percentage levels, where <x> is read off the x-axis. **(D)** same as (C) except matched non-AP triplets were used. **(E)** Model of crowdsourcing effect. ***(a)*** The yellow highlighted region represents a regulatory archipelago (AP) consisting of genes and distal enhancers. Within an AP, spatially proximal binding sites (BS) for a common TF '*crowdsource*' an increase in their own occupancy. Facilitated by increased TF diffusion among large numbers of spatially proximal BS, a spatial homotypic BS cluster favorably alters TF protein concentration in its microenvironment. Predictably, TFs with degenerate motifs, and hence pervasive BS, exhibit the highest occupancy boosts. ***(b)*** In turn, AP enhancers enriched in degenerate motifs experience switch-like multi-fold boosts in accessibility and target gene expression. Overall, a context-specific increase in availability of TFs with degenerate motifs – but not high-specificity motifs – drives a multi-fold boost in chromatin accessibility, thereby underscoring crowdsourcing's likely role in AP activation. ***(c)*** In contrast, a non-AP enhancer does not experience an occupancy boost and activitation. The crowdsourcing mechanism integrates well with the two prevailing models of context-specific gene module activation: in a targeted tissue, higher expression of TFs with a degenerate motif may ***(d)*** induce chromatin loop formation; or alternatively ***(e)*** facilitate release of paused polymerase in pre-formed enhancer-promoter loops. In both cases, crowdsourcing ensures a high degree of context-specificity, mitigating spurious occupancy outside of or AP-active tissue or AP enhancers enriched for degenerate motifs.

### AP activity is correlated with availability of TFs, specifically those with degenerate motifs

Our results thus far suggest that crowdsourcing may be intimately connected to the regulation of AP enhancer-gene complexes, as it provides a way for the cell to prime or induce activity in multiple genomic elements simultaneously, in a specific spatial and tissue context. As shown above, the boost in overall activity (approximated by DHS) of an AP enhancer is in fact far higher in AP enhancers enriched for low-RE (high coverage) BS. However, its is not clear whether the binding of TFs corresponding to the low-RE motifs increases the overall accessibility at the enriched enhancer, or alternatively, an already increased accessibility at enriched enhancer (by some unknown mechanism) is responsible for greater occupancy of particular TFs at those enhancer. In order to resolve this circularity, we tracked tissue-specific gene expression of TFs in 11 cell types and studied its relationship with tissue-specific AP enhancer accessibility (Methods). As shown in Fig. 5C, mean AP enhancer accessibility increases robustly with the expression of TFs comprising high-coverage (red), but not low-coverage, TF-APs (gray), for which there was a slight but significant inverse relationship, also noted in an analysis above (Fig. 2A). No such associations were observed in non-AP enhancer sets controlled for low-and high-RE BS counts (Fig. 5D). Thus, AP enhancer accessibility and activity is highly responsive to the levels of high-coverage AP TFs as they vary across tissues.

Together, these results strongly suggest that crowdsourced boosts in TF occupancy, through the context-specific binding of high coverage TFs, may help drive tissue-specific activation of enhancer networks and their target gene complexes.

## Discussion

### Summary

Here, we have shown that a TF’s *in vivo* occupancy at a particular cognate BS is much greater when the BS is in spatial clustered with other homotypic BS (i.e., in an AP) than when it is not. Strikingly, the size of the occupancy boost robustly scales with the number of BS, or relatedly, the number of enhancers, in the archipelago (AP), suggesting, for the first time, that the BS in an AP cooperatively *crowdsource* their own occupancy. To ensure the robustness of our conclusions, we used stringent controls and employed multiple (i) sources for AP enhancers (Malin et al 2013, Sheffield et al 2013), (ii) experimental backgrounds (non-AP enhancers in the AP-active tissue, the same enhancer in AP-inactive tissues), (iii) occupancy scales (per BS, per enhancer), and (iv) types of occupancy data (curated digital footprints, ChIP-Seq. These observations are not adequately explained by current models, however, they closely agree with a standard biophysical model of facilitated TF diffusion that duly accounts for the augmented diffusion of TFs among spatially proximal homotypic BS. Effectively, a collective of homotypic clusters of TF BS (HCTs) cooperatively alter their microenvironment, raising the local concentration of their cognate TF. The accompanying occupancy boost, in turn, contributes to higher overall enhancer activity and putative target gene expression.

### Genomic versus Spatial homotypicity

Our work synthesizes the regulatory roles of HCTs (e.g. Crocker et al 2015), and of stable chromatin structures (e.g. Dowen et al., 2014), by showing that it is precisely the interplay of numerous HCTs mediated by chromatin folding that gives rise to the previously undocumented biophysical effect that we have termed *crowdsourcing.* Enhancer-enhancer interactions have been reported in the context of HOX and globin gene regulation as well as in high-throughput ChIA-PET assays, but their functional nature has remained elusive. A notable exception, spatial clustering of enhancers around an olfactory receptor gene have been associated with removal of repressive H3K9me3 (Markenscoff-Papadimitriou et al. 2014); it is plausible that crowdsourcing is an upstream trigger of this change – through a general remodeling of the local chromatin state, or through increased binding of a TF that mediates chromatin remodeling.

### Higher order impact of crowdsourcing

Unexpectedly, we observed up to two-fold higher occupancy boost and 10-fold greater normalized chromatin accessibility in AP enhancers enriched for degenerate motifs (‘enriched enhancers’) than in depleted enhancers. A likely explanation is the emergence of an aggregate occupancy effect among an enriched enhancer’s frequent degenerate BS, which serves to remodel the local chromatin state. Under inactive conditions – that is, in AP-inactive tissues or outside of APs – we found that enriched enhancers (which inherently tend toward far lower GC content than depleted enhancers) display substantially higher chromatin accessibility compared to depleted enhancers. This is consistent with previous work suggesting that nucleosomes favor unbound, low GC-content sequence, yet are readily displaced by strongly binding pioneer factors, or, as in the case of crowdsourcing, by an aggregate of distinct TFs (Barozzi et al. 2014; Wasson and Hartemink 2009).

In an AP-active tissue, conversely, enriched AP enhancers experience a widespread surge in binding, thereby displacing the nucleosome and boosting occupancy further, in a positive feedback loop (Fig. 5E). Taken together, the markedly divergent accessibility inside versus outside an active AP confer to enriched enhancers switch-like behavior, where their state is determined by their context: harbored in an AP replete with degenerate homotypic BS, their accessibility increases – but only in tissues in which the cognate TFs are available. In light of this highly context-specific activation and the rapid evolutionary gain of BS for degenerate motifs, we suggest that enriched AP enhancers can evolve adaptively relatively free of consequences from spurious binding. This is the first work to highlight the special functional significance of AP enhancers enriched for abundant, low-affinity BS. However, further work is needed to confirm to what extent depleted enhancers, whose neighbor genes are expressed at up to three-fold lower levels than those of similar non-AP enhancers, have a unique, perhaps repressive, role. Interestingly, genes controlling cell identity in stabilized chromatin structures were found accompanied by repressed genes that coded for yet other lineage-specifying regulators (Dowen et al 2014).

Crowdsourcing integrates well with the two prevailing models of coordinated activation of spatially co-localized gene complexes (Fig. 5E), while providing a missing piece of the puzzle. Whether (a) long-range enhancer-gene loops form *de novo* upon (or along with) activation of a gene cluster (Deng et al. 2012), or (b) the loops are pre-formed and are activated by TF availability (Ghavi-Helm et al. 2014), the cell requires TFs to functionally bind and activate elements specifically in a targeted gene cluster. Crowdsourcing of low-affinity BS is well-suited for such targeting, as it can induce specificity through emergent switch-like binding behavior. Interestingly, a recent study showed a strong correlation between pathway-level gene activity and pathway-level spatial proximity across cell types (Karathia, Hannenhalli et al., manuscript under review), suggesting that chromatin structure is intimately connected with gene complex activation, consistent with (a), above. Unlike direct enhancer-gene interaction in the standard model for distal transcriptional regulation, crowdsourcing, interestingly, is not observable at the level of single enhancer-gene interaction, but instead emerges only at higher levels of chromatin organization and co-regulated gene modules.

### Tissue specificity and cooperative binding

We found that crowdsourcing is highly tissue-specific, as high-coverage AP BS exhibit several-fold greater occupancy in AP-active relative to AP-inactive tissues. Such tissue specificity is consistent with the dependence of crowdsourcing on chromatin context and TF availability, where differential TF availability likely acts not only directly but also by influencing higher-order chromatin conformation (Pombo and Dillon 2015). Crowdsourcing endows the cell with a high degree of fine-grained regulatory control, as occupancy boost magnitude is shaped by the collective availability of multiple TFs and conditioned on the chromatin-induced spatial proximity of their cognate sites. Fundamentally, crowdsourcing provides an alternative mechanism of cooperativity to direct cooperative binding of heterodimerizing TFs, an established source of tissue-specificity. Indeed, crowdsourcing acts complementarily to cooperative binding (Supplementary Note 2).

### Differential occupancy as a vehicle for specificity

In contrast to previous work underscoring the functional importance of weak (low occupancy) binding at non-consensus or inherently weak BS that typical ChIP-Seq processing tends to miss due to stringent cutoffs (Tanay 2006; Biggin 2011; Essien et al. 2009), crowdsourcing leverages spatial chromatin context to imbue inherently low-affinity sites with unexpectedly high-occupancy binding. Unidentified crowdsourcing may therefore underlie previous reports linking particular low-affinity sites with context-specific regulation (e.g. Ramos and Barolo 2013), or linking individual HCTs to unusually robust binding, for example in regulatory regions upstream of HOX genes (Crocker et al., 2015). Indeed, occupancy boosts we observed at spatially clustered HCTs were computed with respect to nominally ‘isolated’ HCTs. As shown by Crocker et al (2015), occupancy is more robust where degenerate homotypic sites are located in genomic clusters. HCTs, however, are highly abundant in the genome (Gotea et al. 2010) as well as, by nature, spatiotemporally invariant, which raises a well-known conundrum, viz. how a TF discriminates among a multitude of candidate BS (Stewart and Plotkin 2013; Stewart et al. 2012). In contrast to the static and relatively low specificity of an individual HCT, a large collective of homotypic low-affinity sites can attain high specificity and spatiotemporal responsiveness precisely by their capacity to reconfigure the local TF environment *en masse* – in specific favorable chromatin contexts. Subject to coordinate regulation, a locus is therefore targeted, not through a fixed, individual address on the one-dimensional genome, but as a conditional and collective nexus in the 3-D chromatin topology – a mobile area code to which the motif's short sequence is appended.

### Potential implications for transcription factories, superenhancers

An archipelago, as described here, represents a group of spatially clustered enhancers and their likely target genes, which are often functionally related (Sheffield et al. 2013; Malin et al. 2013). Meeting this same general description are subnuclear compartments known as transcription factories (Edelman and Fraser 2012). Transcription factories have been shown to concentrate resources such as RNA PolII, core components of transcription, as well as some master TF regulators (Schoenfelder et al. 2010b). However, it is unclear precisely how distinct factories achieve specific and differential concentrations of master regulator TFs (Schoenfelder et al. 2010a). Crowdsourcing offers a possible explanation, and is consistent with a speculated role for resident sequences (Schoenfelder et al. 2010a). While it is generally assumed that high concentrations of TFs are critical in recruiting genes and their distal regulatory regions to the factory, our work suggests alternative causality, as supported by formal biophysical simulations. Although not confirmed, our characterization of archipelagos suggests their operational overlap with factories.

Intriguingly, there is also ample overlap between the conditions for crowdsourcing and known features of superenhancers – enhancers 100Kb or longer in length comprising numerous smaller regions, which regulate genes critical to cell identity (Whyte et al. 2013). Recent works have shown that superenhancers are highly tissue-specific and densely occupied by master regulator TFs, which often recognize degenerate motifs (Vahedi et al. 2015; Heinz et al. 2015). Critically, superenhancers also reveal unusually high levels of spatial interaction among their subunits (Heinz et al. 2015). We thus speculate that crowdsourcing may play a role in superenhancer function.

## Methods

### Enhancer clusters ('APs')

In previous work, genomically dispersed clusters of enhancers with correlated activity across cell lines showed evidence of spatial proximity, particularly in tissues in which the enhancers were active, where spatial proximity between two genomic segments was inferred from Hi-C (Malin et al. 2013). Starting with previously published 40 enhancer clusters, we iteratively filtered out the enhancers from each cluster whose mean spatial proximity to other enhancers was at least one standard deviation below the original mean across all enhancers in the cluster. This results in 40 APs with a total of 1480 enhancers (Supplementary File 1) with ∼37 per AP, ranging from 6 to 89 enhancers per AP.

### Determining *in vivo* occupancy at a BS using digital footprint data

Putative BS in each enhancer were identified using TRANSFAC vertebrate motifs (Matys et al. 2006) and motif scanning tool PWM_SCAN (Levy and Hannenhalli 2002) at 95 percentile score cutoff. We identified *in vivo* TF occupancy by overlapping putative BS with the high-confidence genome-wide digital DNase hypersensitivity footprints identified in 38 human cell lines (Neph et al. 2012). Digital footprints are a single-base-pair resolution readout in which the absence of aligned reads in a particular segment of open chromatin has been shown to predict binding of a protein (Neph et al. 2012). For a TF, a particular putative BS was considered bound by the cognate TF if there was specific overlap between the BS and a footprint, with further requirement that (i) the midpoint of a footprint must overlap the BS; (ii) the midpoint of the BS must overlap the footprint; and (iii) BS length + 1 > footprint length > BS length -4. The latter criteria excludes otherwise significant footprints that are either too short or too long to confidently be associated with a given motif instance. When a footprint strongly overlaps sites for multiple TFs, it was included in the analysis for all such TFs. These highly stringent criteria were applied equally to AP and to non-AP data.

### AP-active and AP-inactive cell lines

For each AP, we identified the cell line in which it was most active. Cell lines deemed active for a given AP are those in which at least 80% of the AP's enhancers are in open chromatin regions, based on overlap with DHS narrow peaks. In case of more than one such tissue, except where noted, we selected the tissue with the highest percentage of open enhancers (see Fig. 1A), depicting workflow. Approximately 95 percent of AP enhancers were found to be accessible in an AP's 'most active tissue', which for the 40 APs, span 15 distinct cell types out of 34 tested.

### Establishing non-AP control

To establish a non-AP control, for each combination of TF and AP enhancer we identified a non-AP enhancer (sampled with replacement) with an identical motif profile, *i.e.* the vector containing the number of instances of each motif mapping to the given TF. This is an important control, as the number of homotypic BS in an enhancer that are cognate to a given TF impacts occupancy (He et al. 2012a). We note that AP and non-AP enhancer have very similar distributions of total BS and length. Additionally, for each TF motif and AP, AP enhancers’ mean DHS in the AP's most active tissue was matched to within 5% in the corresponding non-AP enhancers’ mean DHS in the same tissue. Any TF-AP enhancer pair for which a non-AP could not be found meeting these tight controls was excluded. This procedure yielded 430K AP and non-AP TF-enhancer pairs that harbored 730K BS, of which 31K BS had a DNase footprint suggestive of a binding event.

### Determining TF occupancy at enhancer resolution with ChIP-Seq data

We used ENCODE NarrowPeak Chip-Seq data for 11 cell types – AG10803, AoAF, HA-h, HAEpiC, HCM, HEEpiC, HFF, HIPEpiC, HMF, HMVEC-dBl-Ad, HMVEC-dLy-Neo, HVMF, NH-A, NHDF-Ad, NHLF – which gave a total of 135 TFs and 294 TF-cell line pairs. To increase sample size, all AP-active cell types were used for each AP – that is, all cell types for which 90% of the AP’s enhancers were DNase hypersensitive (Using a cutoff of, alternatively, 80% or 100% did not change the observed trend). Enhancer occupancy by a given TF was determined based on overlap between a <u>+/-</u>50bp window surrounding the ChIP-Seq peak and one or more putative motif instances detected within the enhancer. To mitigate concerns over systematic biases stemming from variability in protocols or labs of origin, we note that all ChIP-Seq data had identical *de facto* weighting for AP and non-AP enhancer-TF pairs, since these were matched by motif for BS counts.

### Comparing neighbor gene expression between AP and non-AP enhancers

As a proxy for an enhancer's target gene, following convention, we used the gene closest to the enhancer. As an extra measure of stringency, in case of non-AP enhancer, we excluded those enhancers that were farther than 50kb from the nearest gene promoter. We paired each AP enhancer with one of the remaining non-AP enhancers while controlling for DHS peak height (within 2%) and numbers of both low-RE and high-RE sites (within 2%), where RE class is based on a (variable) RE threshold. This yielded ∼1200 pairs of AP and matched non-AP enhancers; the exact number varied with the RE threshold. For gene expression, five cell types were used for which overall AP activity, calculated as described above, was at or near its maximum as observed in 15 cell types for which we had digital DNase footprint and RNA-Seq data (www.encodeproject.org/ENCODE). These were HSMM, A549, NHLF, Ag04450, and Bj.

### Estimating TF’s RE

The relative entropy was calculated for each TF motif (i.e., position weight matrix) using TRANSFAC (version 2014.3) (Hannenhalli 2008). In cases where there were multiple motifs associated with a particular TF (coming from different publications etc.), the motif with the lowest RE was chosen, because it is expected to numerically dominate the genome-wide BS for the TF, given its higher degeneracy.

### Identifying low-RE enriched and low-RE depleted AP enhancers

A relative entropy (RE) cutoff was used to classify each putative BS in an enhancer as either low-RE or high-RE (complement of low-RE). This cutoff was varied from RE = 4 (classifies ∼2% enhancers as low-RE) to RE = 9 (classifies > 50% enhancers as low-RE). For each AP enhancer, after tallying the number of low-RE and high-RE sites, an enrichment p-value was generated by applying a Fisher Exact test comparing the numbers of low/high RE sites in the enhancer to those in the pooled set of control (non-AP) enhancers. Based on this enrichment p-value, enhancers were sorted, and the top and bottom ranked x% of enhancers compared in subsequent analysis (x in {10, 20, 50}).

### Calculating a normalized conservation score

To compare evolutionary conservation of low-RE BS enriched AP enhancers to low-RE depleted AP enhancers, we used PhastCons scores, based on 20 mammalian species (Siepel et al. 2005), which are resolved to the individual base. Mean scores across the two classes of enhancers were normalized with respect to non-AP enhancers matched one-to-one with an AP enhancer having approximately the same number of low-RE and high-RE BS, based on a variable RE cutoff, Additionally, we ensured that non-AP enhancers were within 50Kb of the promoter of a highly expressed gene, i.e. its fpkm > 1.0 – which, depending on the cell type, includes approximately the ten percent most highly expressed genes.

### Determining occupancy boost with alternative set of AP enhancers

We obtained sets of correlated regions generated in (Sheffield et al 2013). Each Sheffield cluster of DNase hypersensitive (HS) regions initially spanned multiple chromosomes. To make them consistent with enhancer clusters from Malin et al (2013), regions from a single Sheffield cluster located on distinct chromosomes were treated as distinct clusters, and we retained at most the two largest such clusters from each Sheffield cluster. Consistent with previous procedures, we derived enhancer clusters from each Sheffield cluster by only retaining the regions that overlapped a putative enhancer represented by a large pooled set of 98,000 P300 ChIP-Seq peaks used previously (Malin et al 2013).

To further cull the thousands of resulting enhancer clusters, we excluded those with < 10 enhancers or with mean enhancer DHS < 100 in their most active tissue. We further excluded Sheffield clusters in which fewer than 90% of enhancers were DNase hypersensitive in their most active tissue, resulting in 474 clusters – averaging ∼16 enhancers each, though ranging to over 100. Similar to above (see ‘AP Enhancers’) we used Hi-C data to screen enhancers in each AP that were less spatially proximal, on average, to the remaining members. To prevent excessive removal of additional enhancers, given the already modest mean pre-screen AP size, we implemented the Hi-C screen in a single pass, without recursively updating each enhancer’s mean Hi-C score after removal of a fellow AP member. This resulted in 472 non-empty APs with an average of ∼15 enhancers each.

For background control, we used the complement of P300 ChIP-Seq peaks overlapping any of the screened set of approximately 2.6M Sheffield et al DNase hypersensitive regions. This resulted in too few putative enhancers, and so to this we added back ChIP-Seq peaks overlapping any cluster (on one chromosome) of hypersensitive regions with fewer than five members and with mean DHS > 50 in its most active cell type; this produced a background pool of ∼18K enhancers. Non-AP enhancers from this set were matched with AP enhancers as described above. In order to accommodate the smaller APs in this alternative dataset, we loosened the stringency on DHS control such that mean DHS was matched to within 1% for AP and non-AP sets, overall, with mean DHS for individual TF-AP combinations constrained to less than double or more than half the mean DHS of their non-AP counterpart.

### TF expression-AP activity correlation

This analysis used data from each of 15 cell lines for every AP, encompassing ∼2.4 million BS. Each (TF, AP enhancer, cell line) triplet was assigned (i) a DHS value, corresponding to AP enhancer and cell line; (ii) a coverage score, corresponding to AP enhancer and TF; (iii) a normalized RNA-Seq value corresponding to TF and cell line. Analysis was limited to triplets with a coverage score in the top and bottom 20%. In each of these coverage classes, triplets were further sorted based on the TF’s expression in the given cell line and screened to include only triplets with top or, alternatively, bottom 20 (or 25 or 50) percent TF expression. For each coverage class, the percentage difference in mean cell-type specific DHS between the low TF expression and high TF expression cohorts was plotted. Confidence intervals for each percentage difference were computed on the basis of 50K bootstrap replicates.

### H3K27Ac levels

We downloaded Encode ChIP-Seq peaks from human umbilical vein cells (HUVEC) for histone mark H3K27Ac, known to be associated with active enhancer states (Calo and Wysocka 2013). This cell line was chosen for its combination of available data and a large number of enhancers in APs that are active in the cell line. We compared the ratio in mean ChIP-Seq levels between top 10% enriched and top 10% depleted AP enhancers to the same ratio for non-AP enhancers, matched one-to-one with the AP enhancers as described above. An AP enhancer and its matched non-AP enhancer were included only if the AP enhancer belonged to an AP that was ‘active’ in HUVEC (>80% of its enhancers was DNase hypersensitive). This resulted in ∼40 enriched and ∼100 depleted AP enhancers, and the same number of non-AP enhancers.

### Supplemental information includes

Facilitated Diffusion Model

Extended Experimental Procedures

1 Supplemental Data File

5 Supplemental Figures and Legends

## Acknowledgements

JM wishes to thank Ivan Ovcharenko for valuable feedback throughout the process, and Avinash Das, Kun Wang, and Mahfuza Sharmin for technical assistance.

### Author Contributions

Designed analysis: JM with help from SH and SM. Performed analysis JM with help from HK. Biophysical modelling and simulations: DE, XM. Crowdsourcing mechanism and functional implications: JM. Wrote paper: JM, SH, DE. All authors reviewed paper. SGP helped with illustrations.

